# Exploring Hippocampal Vulnerability: Diminished Angiogenic Capacity in the Hippocampus Compared to the Cortex

**DOI:** 10.1101/2025.02.25.640033

**Authors:** Xiaojie Li, Nanjing Li, Yao He, Qixing Zhong, Zhihui Zhong

## Abstract

**Background:** The hippocampus is one of the first regions affected in various neurodegenerative diseases. In this study, investigated the vascular factors contributing to its susceptibility, aiming to elucidate the underlying vascular mechanisms.

**Method:** Utilizing publicly available single-cell databases, we analyzed the differential expression of genes in blood-brain barrier (BBB)-associated cells within the hippocampus and compared them to those in the cortex. Those genes were further validated in mouse and ischemia rat models.

**Results:** We identified differentially expressed genes (DEGs) in endothelial cells, pericytes, and astrocytes in the BBB. Subsequent gene ontology (GO) enrichment analysis and protein-protein interaction (PPI) network analysis identified key hub genes: *Kdr*, *Fn1*, *Pecam1*, *Cd34*, and *Cd93,* that related to angiogenesis. They differential expression was then experimentally verified using micro-vessels from mouse and rat brains: In the rat ischemia model, we observed up-regulation of angiogenesis-related genes, including *Kdr*, *Cd34*, and *Cd93*, in the microvasculature of both the hippocampus and cortex, with relatively lower expression in the hippocampus.

**Conclusion:** These findings suggest that the hippocampus has a reduced angiogenic capacity compared to the cortex, which may contribute to its increased vulnerability to neurological disorders.

## 1. Introduction

In mammals, significant differences exist between the hippocampus and cerebral cortex in terms of anatomy, cellular structure, and function [1]. The medial cortex forms the hippocampal structure known as the paleocortex, while the dorsal cerebral cortex forms the neocortex. The paleocortex represents an evolutionarily older brain structure, while the neocortex emerged later and underwent significant development in vertebrates, ultimately comprising over 96% of the hemispheric cortex [2]. In the paleocortex and archicortex, neurons are generally arranged into 3 to 5 layers, while the neocortex is structured with 6 layers. The functional region of the neocortex spans the entire cortex, encompassing both primary and advanced sensory areas, primary and secondary motor areas, and other interconnected regions, collectively performing a wide range of integrated functions. The hippocampus is composed of three main regions: the dentate gyrus, the hippocampus proper, and the subiculum. Neurons within these regions are interconnected, forming a network that supports critical functions such as learning, memory, spatial navigation, and emotional regulation [3].

The hippocampus, one of the most extensively studied regions of the brain, is highly sensitive to hypoxia and shows early functional impairments in conditions such as Alzheimer’s disease (AD), epilepsy, vascular dementia, and aging [4, 5]. The level of reactive oxygen species-scavenging tetrahydrobiopterin (BH4) in the hippocampus is lower than that in the whole brain, suggesting that the hippocampus may be more vulnerable to oxidative stress [6]. The interaction between the vascular and nervous systems offers a novel perspective for investigating hippocampal vulnerability [7, 8]. The normal function of neurons is maintained by the blood-brain barrier (BBB), which is primarily composed of brain microvascular endothelial cells, astrocytes, and pericytes. The BBB protects the central nervous system from harmful substances in the blood, regulates the transport of molecules, and helps maintain the stability of the brain’s internal environment [9]. The brain vascular structural and functional impairments are often early indicators of neurological dysfunction [10]. In AD, BBB dysfunction and insufficient cerebral perfusion promote Aβ accumulation and neurofibril entanglement [11]. In neurodegenerative and vascular diseases, the hippocampal BBB is more vulnerable than in the cerebral cortex. Previous studies in our group also confirmed the vulnerability of the BBB in the hippocampus: One year after stroke, the permeability of BBB in the hippocampus of rhesus monkeys was significantly higher than that in the cortex, suggesting that the damage to the BBB in the hippocampus was more severe compared to the frontal cortex [12]; Similar phenomenon was also found in rhesus monkeys with chronic cerebral ischemia. In addition, in the brain-gut axis-related studies of our group, we demonstrated a continuous increase in BBB permeability in the hippocampus of rhesus monkeys with intestinal flora disorder [13]. Furthermore, in the mouse model with intestinal flora disorder, we found that the permeability of BBB in the hippocampus was significantly higher than that in the cortex. Lastly, other studies also demonstrated that the BBB in the hippocampus was more vulnerable than that in the cortex: a study showed that the increase in extracellular histones in the blood is considered a biomarker of vascular dysfunction associated with severe trauma or sepsis, and that histone-induced BBB leakage occurs more frequently in the hippocampus compared to the cortex [14]. In 2021, Shaw. K et al compared the neurovascular function of the hippocampus and visual cortex in conscious mice by measuring two-photon imaging of single neurons and blood vessels, local blood flow, and hemoglobin oxygenation. The study found that, due to the difference in vascular network, pericyte, and endothelial cell function, the blood flow, blood oxygen and neurovascular coupling in the hippocampus were less than that in the neocortex, and these characteristics may limit the supply of oxygen. This also explains its sensitivity to injury in neurological diseases [8].

Therefore, studying the injury mechanisms of BBB in the hippocampus is crucial for maintaining its normal structure and function. We aim to investigate why the BBB in the hippocampus is more vulnerable compared to the cerebral cortex. Understanding the underlying factors contributing to this increased susceptibility could provide valuable insights into neurovascular dysfunction and its impact on hippocampal health. Currently, studies on the heterogeneity of hippocampal and cortical damage primarily focus on the neurons themselves, while research on the vulnerability of the BBB in the hippocampus remains limited. Most reports focused on the phenotypes associated with specific biological processes, while some studies have examined the overall structure and molecular foundations of cerebral vessels. This is crucial for advancing our understanding of cerebral circulation and cerebrovascular diseases. Visualizing cerebral vessels also gives us a direct understanding of the differences in blood vessels across different regions of the brain. In 2020, Miyawaki, T. et al. established a method of three-dimensional (3D) visualization and molecular representation of the whole vascular network named “SeeNet”, which includes almost all the micro-vessels in the whole brain [15], provided a new idea for us to study the vascular heterogeneity of the hippocampus and cortex. In addition, Micro-Optical Sectioning Tomography (MOST) can image the cerebral vessels of mice at the submicron resolution [16]. After comparing the vascular structure of wild-type (WT) mice and transgenic AD mice, the blood vessels of transgenic AD mice seem to be thinner, more uneven, and less organized. Furthermore, single-cell level sequencing and proteomics are essential tools for investigating the underlying mechanisms. These techniques also help to explain the vascular heterogeneity of the BBB in the hippocampus and cortex [2, 17].

In this study, we used single-cell sequencing as our starting point. Single-cell transcriptome sequencing provides valuable insights into cellular heterogeneity that cannot be detected by bulk sample sequencing. This approach allows us to analyze the different cellular components of the BBB. By mining publicly available cerebral vascular-related single-cell sequencing databases, we were able to identify and analyze the differential genes expressed in vascular endothelial cells from the hippocampus and cortex of the brain.

## 2. Materials and methods

### 2.1 Single-cell RNA sequencing (scRNA-seq) database

In this study, we employed relevant datasets obtained from the GEO database (http://www.ncbi.nlm.nih.gov), including the mouse datasets GSE185862 [18], SRP135960 [19], GSE60361 [20], and the human dataset [21], and performed quality control (QC) to assess mitochondrial gene content, gene count, and other indicators to identify and exclude potential multi-cell or doublet contamination. After QC, the data were imported into Seurat objects. Subsequently, we applied the Seurat 4.0 package for data normalization, dimensionality reduction, and clustering analysis, to remove batch effects and obtain reliable data. Using the FindClusters function, we clustered the cells into distinct groups based on the reduced dimensions. Cell type annotation was carried out by referencing known cell markers from the literature, as well as markers commonly used in similar studies. Additionally, the machine learning tool scType was employed to automate the annotation of cell types for each cluster. The data also included tissue region information, such as cortical and hippocampal regions, providing a contextual background for the analysis.

### 2.2 Identification of differentially expressed genes (DEGs)

Differential gene expression analysis was conducted using the Seurat’s FindMarkers function in Seruat4.0 by setting reference and comparison groups (cortex vs hippocampus). Ggplot2 was used to construct volcano maps to visualize the DEGs. Significant differentially expressed genes were identified using thresholds of log_2_FC > 0.25 and *P* < 0.05.

### 2.3 Functional and pathway enrichment analysis

Gene Ontology (GO) analysis was used to identify the characteristic biological attributes of genes, gene products, and sequences, including biological process (BP), cell components (CC), and molecular function (MF). Enrichment databases of human and mouse from ENSEMBLE GRCh38.p13, GRCm39, and GO were used. Using R clusterProfiler 4.4.4, the DEGs we used in the previous step were functionally enriched. BiomaRt_2.52.0 records the terms of some functionally associated databases.

### 2.4 PPI network construction and hub genes identification

We employed an online tool (http://www.string-db.org/) for identifying and predicting interactions between genes or proteins and constructed the PPI network of intersecting DEGs. Next, we used the Cytoscape software to visualize the DEG’s PPI network. The key nodes in the network were investigated using the Cytoscape plug-in Cytohubba to explore the central genes contained in the PPI network.

### 2.5 Isolation of cerebral micro-vessels

The micro-vessels of the hippocampus and cortex were extracted by density gradient centrifugation following the reported protocol [22], and the targeted genes were screened and analyzed using real-time quantitative PCR (RT-qPCR). After sacrificing the animals (mice and rats), their brains were removed and the cortex and hippocampus were isolated immediately. The brain tissue was homogenized, centrifuged and the supernatant was removed. 15% Dextran 70 (Sangon, 9004-54-0, China) was added and centrifuged again to separate the upper and lower layers, with the vascular pellet remaining in the lower-most layer. The pellet was re-suspended in PBS and filtered through the 40 μm cell sieve. The microvascular segment was left in the upper layer of the filter. After inverting the filter and recovering with the culture medium and then centrifuging, the pure microvascular precipitate was obtained.

### 2.6. Two-vessel occlusion (2VO) modelling

We established a rodent model of chronic cerebral blood flow hypoperfusion-induced pathophysiological changes and cognitive impairment using bilateral common carotid artery ligation in rats (termed the 2VO rat model). This model effectively simulates the conditions of hypoperfusion in the brain. The 2VO model rat is achieved by ligating bilateral common carotid arteries. Ligation of bilateral common carotid arteries in rats can reduce the cerebral blood flow and simulate the pathological changes caused by chronic ischemia, thus, it is widely used in the study of chronic cerebral ischemia [23]. Rats for modelling were prepared using male Wistar rats aged 9–12 weeks (weight, 300–350 g). Under anesthesia with 1–3% halothane inhalation, the bilateral common carotid arteries (CCA) of the rats were exposed through a midline cervical incision and permanently ligated. After carefully incising the carotid artery sheath and separating the vagus nerve from the CCAs, only the CCAs were ligated. During the procedure, the body temperature of the rats was maintained using a heating pad.

### 2.7 RT-qPCR

Total RNA was extracted from the cell samples using the TRIzol™ Reagent (Invitrogen, 15596026, American) according to the manufacturer’s instructions. Reverse transcription was performed using the HiScript III RT SuperMix for qPCR (Vazyme, R323-01, China). The synthesized cDNA was stored at -80[ until further use. RT-qPCR was conducted using the ChamQ Universal SYBR qPCR Master Mix (Vazyme, Q711-02, China). The relative expression levels of the target genes were calculated using the 2^(-ΔΔCt) method, with β-actin (*Actb*) as the internal reference gene. The PCR primers used in this study were listed in **Table 1** below.

**Table 1.**
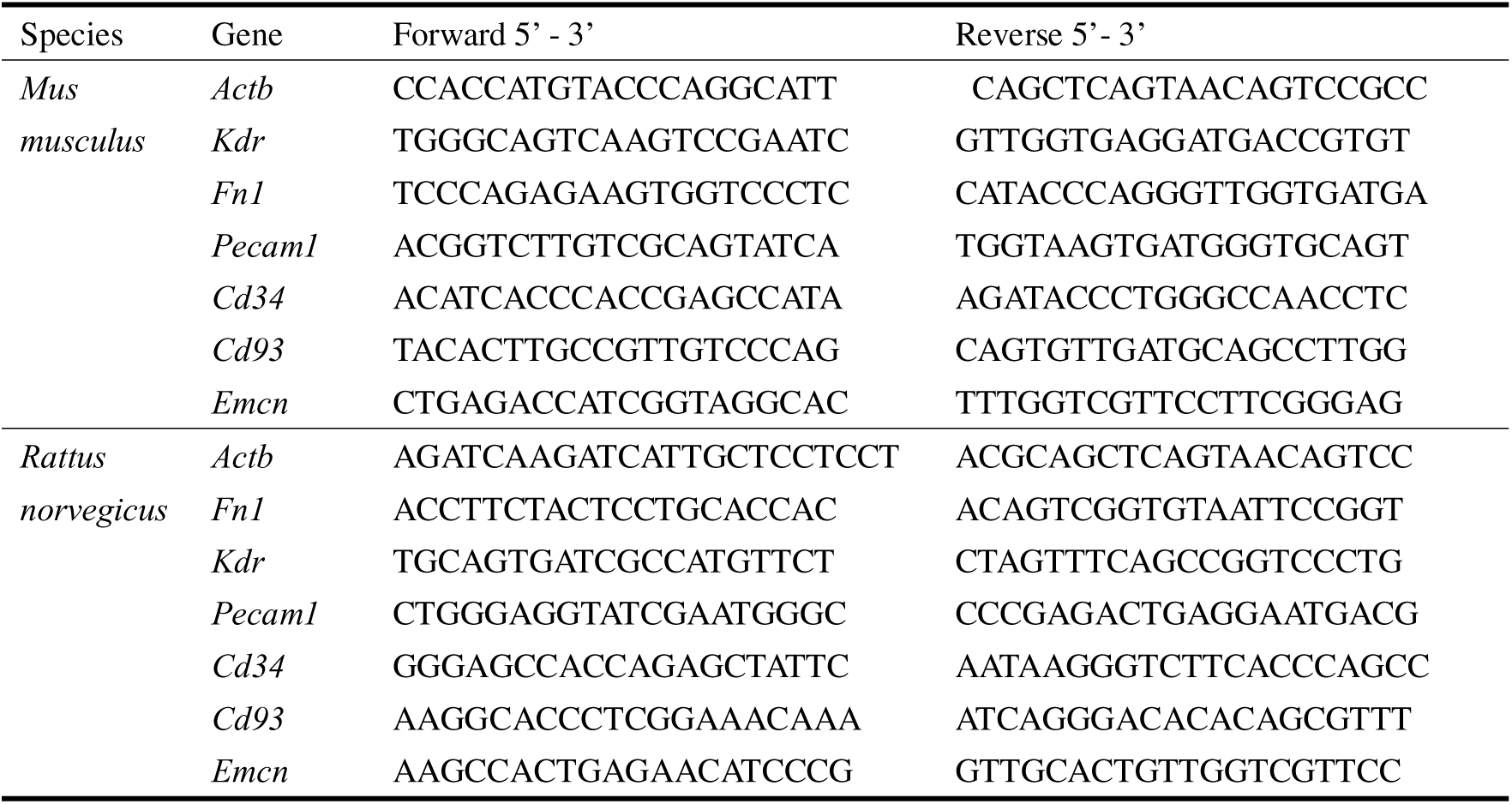
Sequence of the PCR primers.

### 2.8 Statistics analysis

Graphs and statistical analyses were performed using Prism 8 software (GraphPad). Comparisons among multiple groups were conducted using two-way ANOVA, while a two-tailed unpaired Student’s t-test was applied for comparisons between two groups. Data were presented as mean±standard error of the mean (SEM), with a *P* < 0.05 considered statistically significant.

## 3. Results

### 3.1. DEGs between the cortical and hippocampal BBB were identified

To explore the differences of BBB between the cerebral cortex and hippocampal regions in both the mouse and human brain, we investigated three cellular components of the BBB: endothelial cells, pericytes, and astrocytes. We then compared the DEGs of these three types of cells in the cerebral cortex and the hippocampal region, respectively, and the DEGs were presented with volcano plots. The general workflow of this study is shown in **Fig.1** below.

**FIGURE 1.**
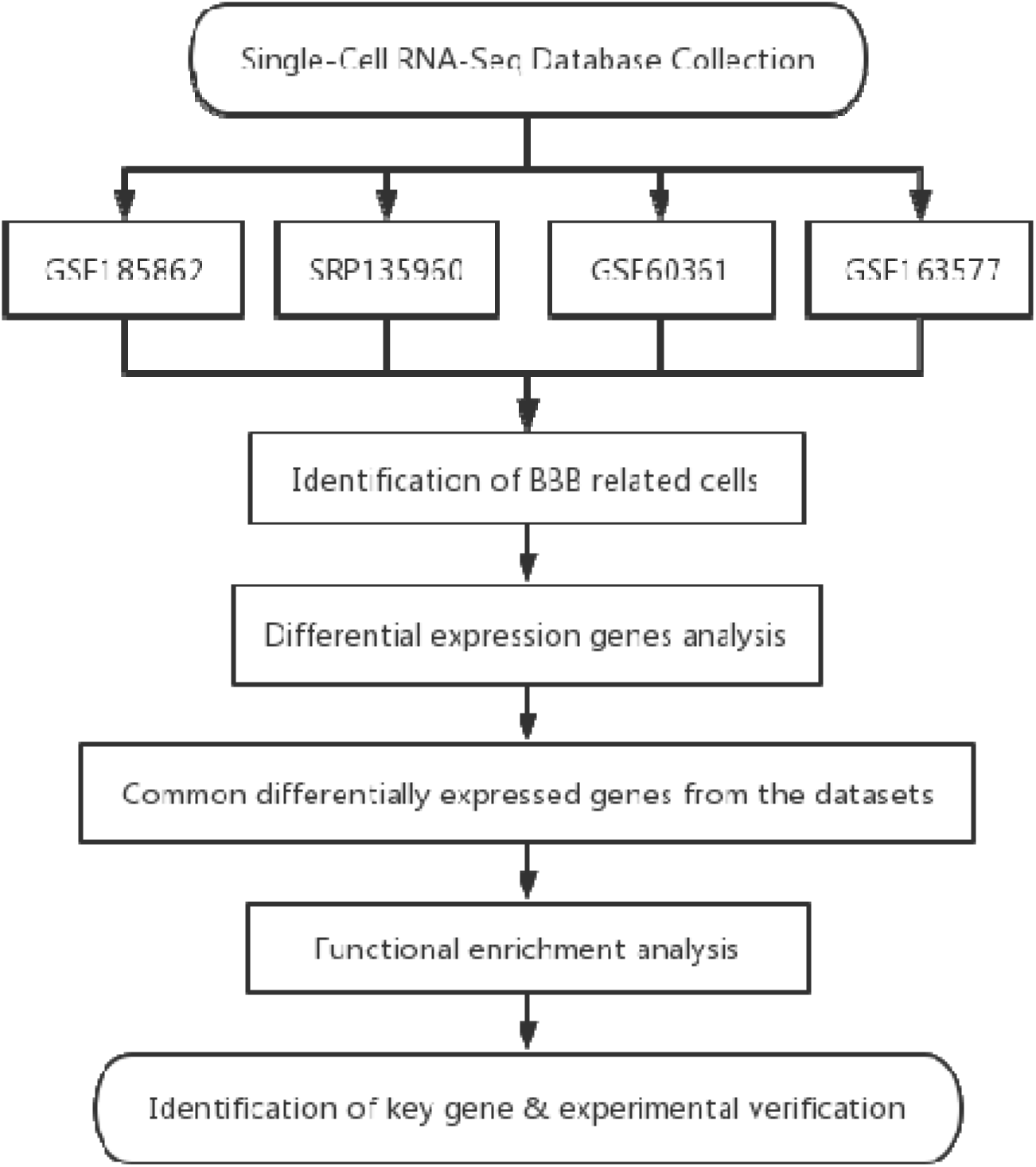
Flowchart of this study. We selected four datasets for our analysis: the mouse datasets GSE185862, SRP135960, GSE60361, and the human dataset GSE163577. From these datasets, we extracted three key cellular components of BBB: endothelial cells, pericytes, and astrocytes. We then compared the DEGs of these cell types between the cerebral cortex and hippocampus of the brain, followed by functional enrichment analysis of the identified DEGs and the experimental validation.

### 3.2 Angiogenesis-related genes were more highly expressed in cortical endothelial cells than in those of the hippocampus

To investigate the disparities in BBB characteristics between the hippocampus and the cortex, as well as to elucidate the susceptibility of the hippocampus, we conducted a comparative analysis of differential gene expression in the endothelial cells, which are the major components of the BBB, in the mouse and human brain: GSE185862, SRP135960, GSE60361, and GSE163577, respectively. The volcano plots of the highly or lowly expressed genes were shown in **Fig. 2A-D**. We identified that the expressions of genes *Bsg, Id1*, *Cxcl12* (**Fig. 2A**), *Tm4sf1*, *Calm2* (**Fig. 2B**)*, Gpx4*, *Cst3, Sema3e* (**Fig. 2C**), *EGFL7*, *NEAT1*, and *PTH1R* (**Fig. 2D**), which were related to angiogenesis, were higher in cortical endothelial cells than in the hippocampus. Moreover, the expression levels of inflammation-related genes *Jun*, *Junb* (**Fig. 2B**), and *IFITM3* (**Fig. 2D**) in cortical endothelial cells were lower than those in the hippocampus. Such inflammatory signaling may suppress neurogenesis in the hippocampus, which may partially explain the vulnerability of the hippocampus [4, 21]. After the GO enrichment, we discovered that genes highly expressed in cortical endothelial cells, as compared to those in the hippocampus, were enriched in pathways directly or indirectly related to angiogenesis, such as the cell-cell adhesion, positive regulation of angiogenesis, angiogenesis, negative regulation of extracellular matrix disassembly, response to oxidative stress, and vascular endothelial growth factor receptor signaling pathway (**Fig. 2E-H**, related DEGs were listed in **Table 2**). The pathways enriched with genes that were lowly expressed in the cortex endothelial cells compared to the hippocampus were shown in **Fig. 2I-L**, only the pathway “negative regulation of angiogenesis” was identified to be associated with angiogenesis. Collectively, our findings suggest that, compared to the cortex, the hippocampus may have a limited angiogenic capacity, making it less equipped to cope with damage caused by various diseases.

**FIGURE 2.**
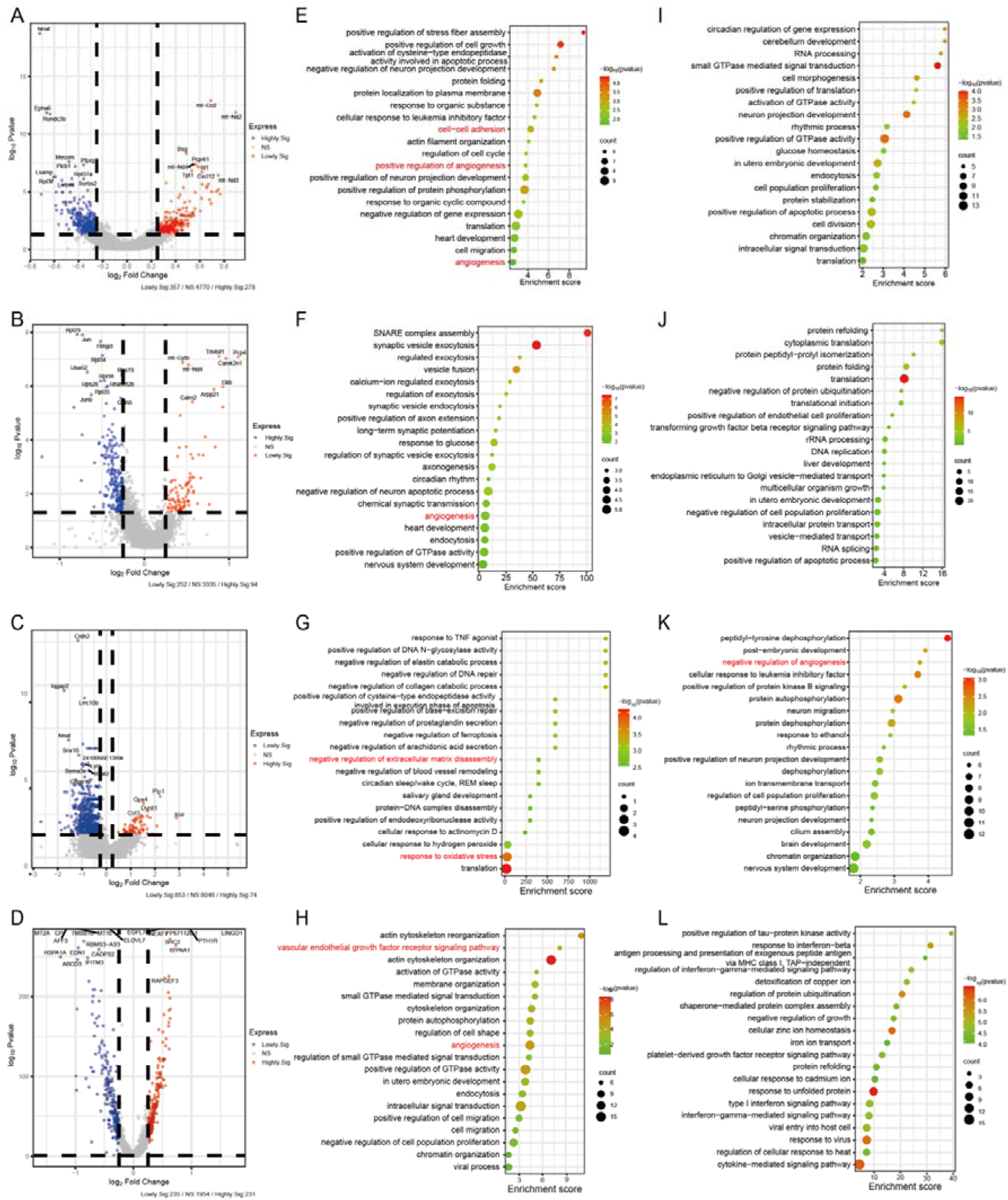
Identification of DEGs and functional enrichment in the endothelial cells (cortex VS hippocampus). (**A-D**) Volcano plots of DEGs (cortex VS hippocampus) for GSE185862, SRP135960, GSE60361, and GSE163577 databases, respectively. The following items were presented in the same order. Data points in red represent highly expressed genes while the blue represent lowly expressed genes. The differences were set as |log FC|>0.25, *P* < 0.05. (**E-H**) GO enrichment analysis of highly expressed genes. (**I-L**) GO enrichment analysis of lowly expressed genes.

**Table 2.**
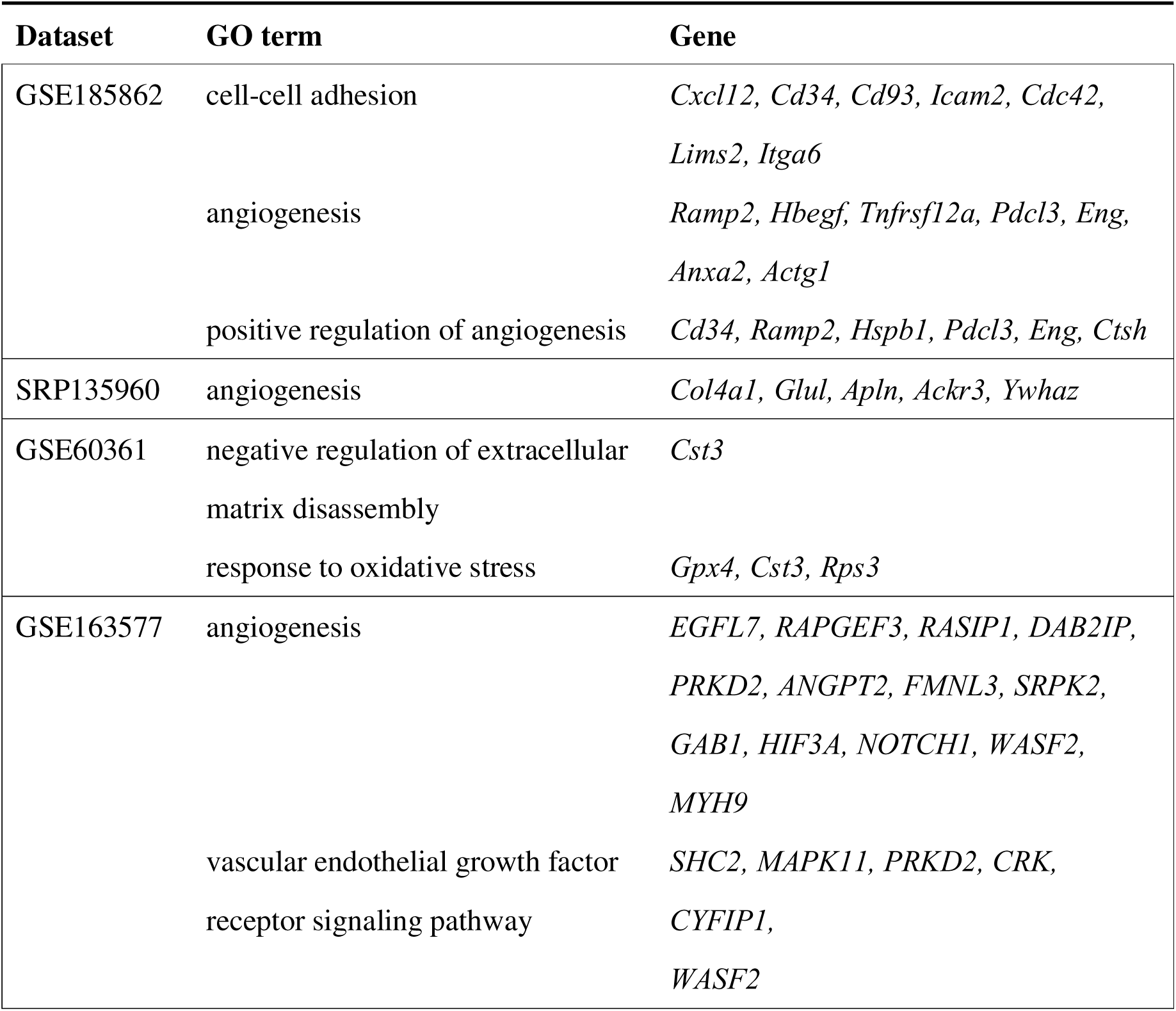
DEGs identified that related to angiogenesis in the endothelial cells (cortex VS hippocampus).

Pathways identified to be associated with angiogenesis were highlighted in red color. The counts represented the number of genes enriched in the corresponding pathway, with larger dots indicating a higher number of enriched genes. The different colors corresponded to varying -log_10_(p-value), with values closer to red indicating smaller p-values, and those closer to green representing larger p-values.

### 3.3 Angiogenesis-related genes were more highly expressed in cortical BBB cells than in those of the hippocampus in mice

To further investigate the above conclusion, we continued to analyze the mouse brain micro-vessels using the database. Considering that the extracted brain micro-vessels include the three cellular components of the BBB: endothelial cells, astrocytes, and pericytes [22], we further integrated these cells into a single cell population as“BBB cells” in three mouse datasets (GSE185862, SRP135960, GSE60361) and examined their differential gene expression between the cortex and hippocampal regions. The results showed that, compared to the hippocampus, 94 genes were commonly lowly expressed in the cortex (**Fig. 3A**), with their GO analysis presented in **Fig. 3C**. Conversely, 62 genes were commonly highly expressed (**Fig. 3B**), with their GO analysis shown in **Fig. 3D**. Specifically, compared to the hippocampus, although none of the pathway was found associated with angiogenesis in the lowly expressed genes (**Fig. 3C**), pathways related to angiogenesis were enriched in the highly expressed genes in the cortex (**Fig. 3D**, highlighted in red). These results further confirmed that the hippocampus has a reduced capacity for angiogenesis compared to the cortex. The protein-protein interaction (PPI) network was then constructed to visualize the interactions between proteins encoded by angiogenesis-related genes. The size and color of the nodes represented their relative significance or connectivity within the network (**Fig. 3E**). For instance, *Kdr* serves as the central hub with the highest degree of connectivity, highlighting its critical role in this network. From the results, several hub genes were identified, such as *Kdr, Fn1*, and *Pecam1*, that are known to play essential roles in angiogenesis which is essential for the repair of vascular and neural injuries [24-26]. Interestingly, among these hub genes in BBB cells, *Cd34*, *Cd93*, and *Cxcl12* were also highly expressed in cortical endothelial cells (cortex vs. hippocampus, **Table 2**). This observation underscored the reliability and robustness of our predicted hub gene results,

**FIGURE 3.**
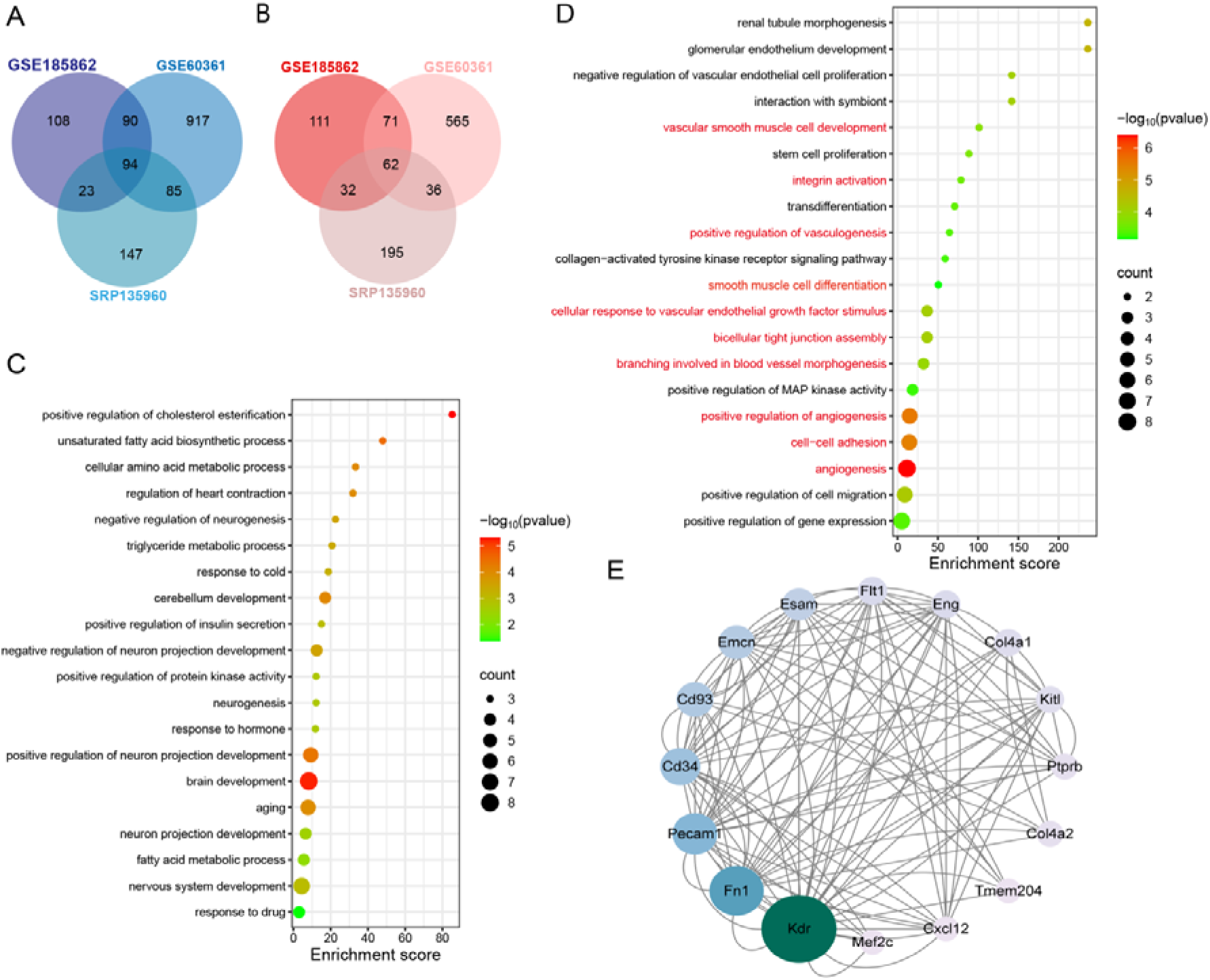
Common DEGs analysis and functional enrichment in BBB cells (cortex VS hippocampus). Venn diagram of common DEGs in a combined set of endothelial cells, astrocytes, and pericytes across three mouse datasets (GSE185862, SRP135960 and GSE60361). **(A)** 94 common DEGs were identified to be lowly expressed in the three datasets. **(B)** 62 common DEGs were identified to be highly expressed in the three datasets. **(C)** GO enrichment analysis of commonly lowly expressed DEGs. **(D)** GO enrichment analysis of commonly highly expressed DEGs. The counts represented the number of genes enriched in the corresponding pathway, with larger dots indicating a higher number of enriched genes. Different colors corresponded to varying -log_10_(p-value), with values closer to red indicating smaller p-values, and those closer to green representing larger p-values. The pathways highlighted in red color are associated with angiogenesis. **(E)** PPI network of highly expressed genes associated with angiogenesis. The size of each dot represents its betweenness centrality.

### 3.4 Angiogenic capacity is lower in the hippocampus than in the cortex of the mouse brain

To further validate our hypothesis that the hippocampus has a lower angiogenic capacity than the cortex, we isolated brain micro-vessels from both the cortex and hippocampus of naïve mice, respectively, to investigate the expression changes of the top 6 hub genes that identified to be associated with the angiogenesis (shown in **Fig. 3E**). Our results demonstrated that, compared to the hippocampus, the genes *Fn1* and *Pecam1* were significantly highly expressed in the cortical micro-vessels (*P* < 0.05), while the genes *Kdr*, *Cd34*, and *Cd93* showed a notable trend towards higher expression (**Fig. 4**).

**FIGURE 4.**
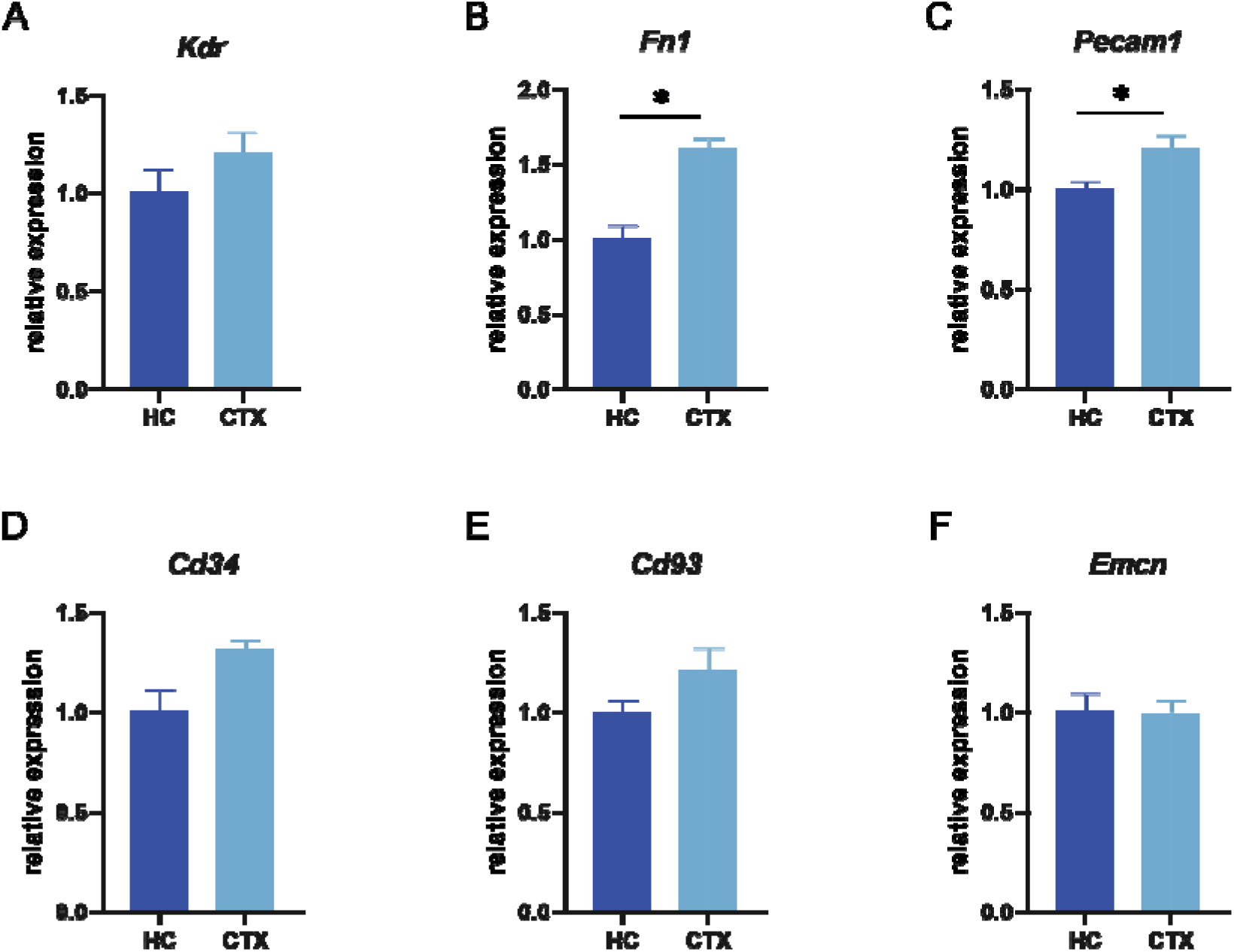
Validation of hub genes using mouse micro-vessels. The relative mRNA expression levels of target genes were quantified using RT-qPCR and normalized to the housekeeping gene *Actb.* Data were presented as fold changes relative to the hippocampus. HC: hippocampus, CTX: cortex. (**A**) *Kdr*, (**B**) *Fn1*, (**C**) *Pecam1*, (**D**) *Cd34,* (**E**) *Cd93*, (**F**) *Emcn*. Data were presented as mean ± SEM, (n = 3), and asterisks indicate statistical significance (\**P* < 0.05).

### 3.5 Angiogenic capacity is lower in the hippocampus compared to the cortex in rat 2VO model

To further validate our hypothesis, we established a rat model under ischemic and hypoxic conditions. It has been reported that genes associated with angiogenesis were up-regulated in these conditions, potentially leading to the formation of new blood vessels to provide increased oxygen and nutrient support to the ischemic area [27, 28]. In other studies, to induce cerebral hypoperfusion in mice, a bilateral common carotid artery stenosis (BCAS) model was established by partially restricting blood flow with micro-coils placed around the bilateral common carotid arteries, resulting in chronic hypoperfusion [29]. Furthermore, in rats, permanent ligation of the bilateral common carotid arteries rapidly induces cerebral hypoperfusion. Compared to the mouse BCAS model, this 2VO rat model is straightforward, producing a more severe and rapid ischemic condition that can lead to significant global brain damage [30].

We used the same method for dissecting the cerebral cortex and hippocampus and extracting blood vessels in mice to isolate brain micro-vessels from both sham-operated and 2VO rats. From our subsequent RT-qPCR analysis with above mentioned hub genes (shown in **Fig. 3E**), among them, *Kdr*, *Cd34*, and *Cd93*, were up-regulated in the vasculature of both the hippocampus and cortex, but their expression levels were significantly lower in the hippocampus compared to the cortex (**Fig. 5A, D, E**). *Fn1 and Pecam1* showed differences between the hippocampus and cortex only for three days and one day after modelling, respectively (**Fig. 5B, C**), which may be explained as the temporal dynamics of hypoxia-regulated angiogenesis [31]. However, three days after modelling, the expression of the *Emcn* gene in hippocampal micro-vessels was significantly higher than that in the cortex (**Fig. 5F**). This finding suggested the involvement of unrecognized mechanisms related to angiogenesis in the hippocampus, warranting further investigation.

**FIGURE 5.**
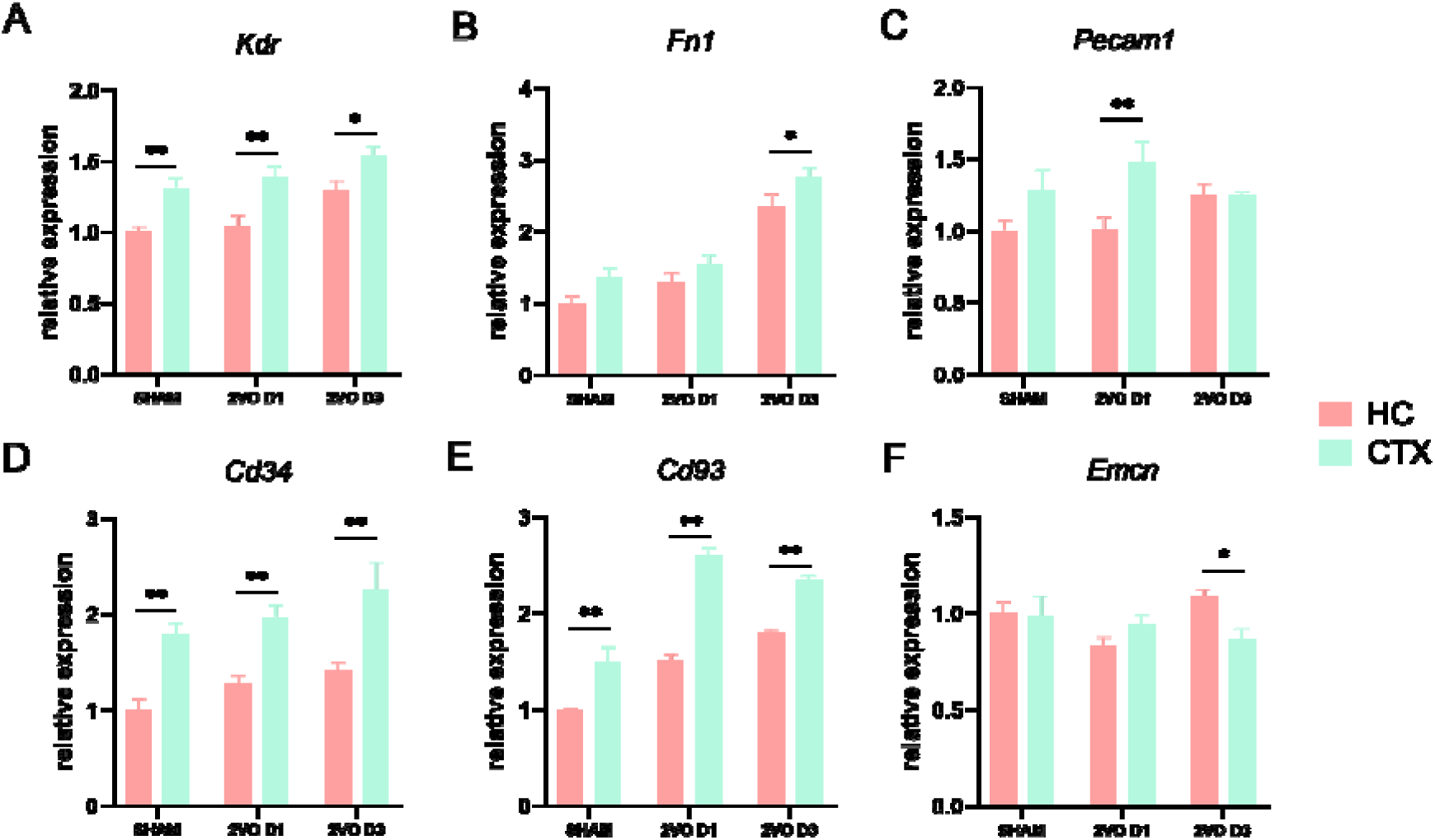
Validation of hub genes using 2VO rat micro-vessels. The relative mRNA expression levels of target genes were analyzed in sham-operation, day 1 and day 3 after modelling, and were quantified using RT-qPCR and normalized to the housekeeping gene *Actb.* Data were presented as fold changes relative to sham-operated rats. HC: hippocampus, CTX: cortex. (**A**) *Kdr*, (**B**) *Fn1*, (**C**) *Pecam1*, (**D**) *Cd34,* (**E**) *Cd93*, (**F**) *Emcn*. Data were presented as mean ± SEM (n = 3). Asterisks indicate statistical significance (\**P* < 0.05, \*\**P* < 0.01).

## Discussion

Angiogenesis plays a pivotal role in the repair of neurovascular injuries and represents a critical process for restoring the function of damaged tissues. In the context of injury, angiogenesis facilitates the formation of new capillary networks, providing ischemic and hypoxic regions with adequate oxygen and nutrient supply, thereby promoting neuronal survival and metabolic recovery [27, 28]. Moreover, newly formed blood vessels offer essential pathways for clearing metabolic waste and inflammatory mediators, contributing to the improvement of the local micro-environment and alleviation of secondary damage [32]. Furthermore, angiogenesis is closely linked to neurogenesis, with both processes mutually enhancing each other through shared signaling pathways, such as vascular endothelial growth factor (VEGF). For instance, the administration of VEGF in central nervous system injuries has been shown to simultaneously promote angiogenesis and neurogenesis, thereby facilitating functional recovery [33]. Newly formed blood vessels not only provide physical support for axonal extension and synapse formation but also secrete growth factors and chemokines that directly stimulate neuronal repair and regeneration [34, 35].

Our study, utilizing bioinformatics analysis, demonstrated that genes highly expressed in the cortex, as compared to the hippocampus, were enriched in angiogenesis-related pathways. This finding suggested that the hippocampus may have a reduced capacity for angiogenesis relative to the cortex. Consequently, the hippocampus may be unable to generate enough new blood vessels for compensatory purposes, which is essential for repairing vascular and neural injuries caused by various diseases.

To validate this hypothesis, we extracted samples from the micro-vessels of mouse brain and experimentally verified the hub genes identified from the bioinformatics analysis of mouse datasets. The results showed that the expression levels of genes *Kdr*, *Fn1*, *Pecam1*, *Cd34,* and *Cd93* were higher in the cortex compared to the hippocampus, which were consistent with the findings from the bioinformatics analysis. In rats, the expression of these hub genes was higher in the cortex than in the hippocampus, consistent with observations in mice, which may be attributed to their genetic homology. Compared to the mouse BCAS model, the rat 2VO model more rapidly induces a more severe state of global cerebral ischemia. Interestingly, in the 2VO ischemic rat model, genes associated with angiogenesis, such as *Kdr*, *Cd34,* and *Cd93*, showed compensatory increases. In addition, the levels of these genes in the hippocampus remained lower than in the cortex, further supporting our hypothesis. Future research should include more disease models and functional validations of the hub genes. Due to the challenges in obtaining human samples, it will also be important to validate the angiogenesis-related genes identified from human databases in suitable models, such as human brain organoid models.

## Conclusion

In conclusion, our study offers valuable insights into the vulnerability of the hippocampus, suggesting that, compared to the cortex, it may have a reduced capacity for angiogenesis. This reduced capacity affected its ability to supply oxygen and nutrients to damaged tissues, clear harmful mediators, and support neuronal survival and neurogenesis, all of which are crucial for brain repair.

## Declarations

## Acknowledgement

The authors would like to thank Sichuan Junhui Biotechnology Co., Ltd. and Singapore HumanSim Pte. Ltd. for providing the experimental platforms for this study.

## Author contribution

The project was conceived and supervised by Z.H. Zhong and Q.X. Zhong. Bioinformatics analysis was conducted by X.J. Li, and N.J. Li. X.J. Li and Y. He carried out the biological experiments and drafted the manuscript. The manuscript was revised by Z.H. Zhong and Q.X. Zhong. All authors approved the final version of the manuscript.

## Availability of data and materials

Data will be made available on reasonable request.

## Ethics approval and consent to participate

Not applicable

## Consent for publication

Not applicable

